# The CREB-regulated co-activators 2/3, have a role, *in vivo*, in osteoblastic gene expression

**DOI:** 10.64898/2026.09.21.753306

**Authors:** Carole Le Henaff, Jobin Joseph, Whitney Petrosky, Zhiming He, Aditya Mohapatra, Sohan Sirimarapu, Jennifer J. Westendorf, Marc Montminy, Nicola C. Partridge

**Author notes:** Corresponding Authors: Nicola C. Partridge, Ph.D., Center for Advanced Biotechnology and Medicine, 679 Hoes Lane West, Room 204, Piscataway, New Jersey 08854-8021; Carole Le Henaff, Ph.D., Center for Advanced Biotechnology and Medicine, 679 Hoes Lane West, Room 132, Piscataway, New Jersey 08854-8021.

## Abstract

Many hormones and substances acting through G-protein coupled receptors and protein kinase A (PKA) activation inhibit the salt-inducible kinases (SIKs) by phosphorylation. SIKs tonically phosphorylate CREB-regulated transcriptional coactivators (CRTC1, 2 and 3), sequestering them in the cytoplasm and, thus, preventing their translocation into the nucleus. Once in the nucleus, CRTCs bind CREB family member transcription factors and enhance their activity. We and others have shown that parathyroid hormone (PTH) activation of PKA and resultant SIK2/3 inhibition allows CRTC2/3 nuclear translocation. One of the major actions of CRTC2/3 in the osteoblast lineage is the regulation of transcription of *Rankl*, as well as other PTH-controlled genes. However, little is known about the role of these co-activators in the osteoblast lineage *in vivo.* Here, we have investigated whether there are basal effects *in vivo* on bone examined at 2 different ages of conditional deletion of these two co-activators in the osteoblast lineage using Col2.3-Cre. We found significant increases in body weight, length, bone mineral density, bone volume/total volume, trabecular thickness and number with decreased trabecular separation in young (2 months old) male mice, all of which dissipated by 6 months of age. Female mice showed minimal changes in the bone phenotype at either age. Nevertheless, there were gene expression changes in bones of both sexes at both ages, and in particular decreases in *Rankl*, *Runx2* and *Sost*, and accompanying changes in Wnt pathway genes. These effects may explain the changes in the bone phenotype in the young male mice, but it is notable that there is a sexual dimorphism in the action of CRTC2 and CRTC3. Overall, the work supports the data from research *in vitro* and forms a basis for investigation of the role of these co-activators in PTH action *in vivo*.

## Introduction

Many hormones and substances acting through G-protein coupled receptors and protein kinase A (PKA) activation inhibit the salt-inducible kinases (SIKs) by phosphorylation. SIKs tonically phosphorylate CREB-regulated transcriptional coactivators (CRTC1, 2 and 3), sequestering them in the cytoplasm^(1)^ and, thus, preventing their translocation into the nucleus. Once in the nucleus, CRTCs bind CREB family member transcription factors as a tetramer, increasing the latter’s action by stabilizing the transcription factors on the DNA^(2)^.

Parathyroid hormone (PTH) is one of the hormones acting in this way on osteoblast lineage cells. We and others have shown, with osteoblast and osteocytic cultures, that PTH activation of PKA and resultant SIK2/3 inhibition, coupled with action of protein phosphatases 1, 2, 4 and 5, causes CRTC2/3 phosphorylation levels to decrease and allows their nuclear translocation^(3-6)^. One of the major actions of CRTC2/3 in the osteoblast lineage is the regulation of transcription of *Rankl*, as well as other PTH-controlled genes^(3, 5-7)^. CRTC1 does not appear to be important in this pathway, while CRTC2 is the key effector for PTH-induced *Rankl* transcription in osteoblasts with assistance from CRTC3, especially the latter in highly differentiated osteoblasts^(3, 5)^. As a result, we have focused in the present report on the roles of CRTC2 and CRTC3 *in vivo*. Single global knockouts of these two genes generated relatively mild metabolic phenotypes^(8, 9)^; yet double global knockouts were embryonic lethal^(10)^. Conditional knockout mice were developed for both genes^(11, 12)^ with a mild phenotype with inducible deletion of CRTC2 in beta cells of the pancreas^(11)^while Prx1-Cre deletion of both *Crtc2* and *3* resulted in increased bone marrow neutrophils and plasma G-CSF levels^(10)^. These authors attributed the changes to effects on bone marrow stromal cells. From the little information provided, it appears there were no gross changes in the sizes of the long bones, suggesting that no major developmental events occurred. In the following work, we have investigated whether there are basal effects *in vivo* on bone of conditional deletion of these two co-activators in the osteoblast lineage using Col2.3-Cre. We have examined bone mineral density (BMD), microarchitecture and gene expression in male and female mice at 2 different ages.

## Results

### Deletion of CRTC2 and CRTC3 in osteoblasts improved the bone microarchitecture in young male mice

*Crtc2* and *Crtc3* were deleted specifically in osteoblasts using a constitutively active type 1 collagen promoter (Col2.3Cre). At 2 months old, male *Crtc2/3^ob-/-^* mice presented increased body weight (Figure 1A) and length (Figure 1B). These mice were followed over time by Dual X-Ray imaging (DEXA-PIXImus). The analyses showed increased bone mineral density (BMD) in femurs (Figure 1D), and a tendency to increase in whole body (Figure 1C), and tibiae (Figure 1E) but without changes in vertebrae (Figure 1F).

**Figure 1:**
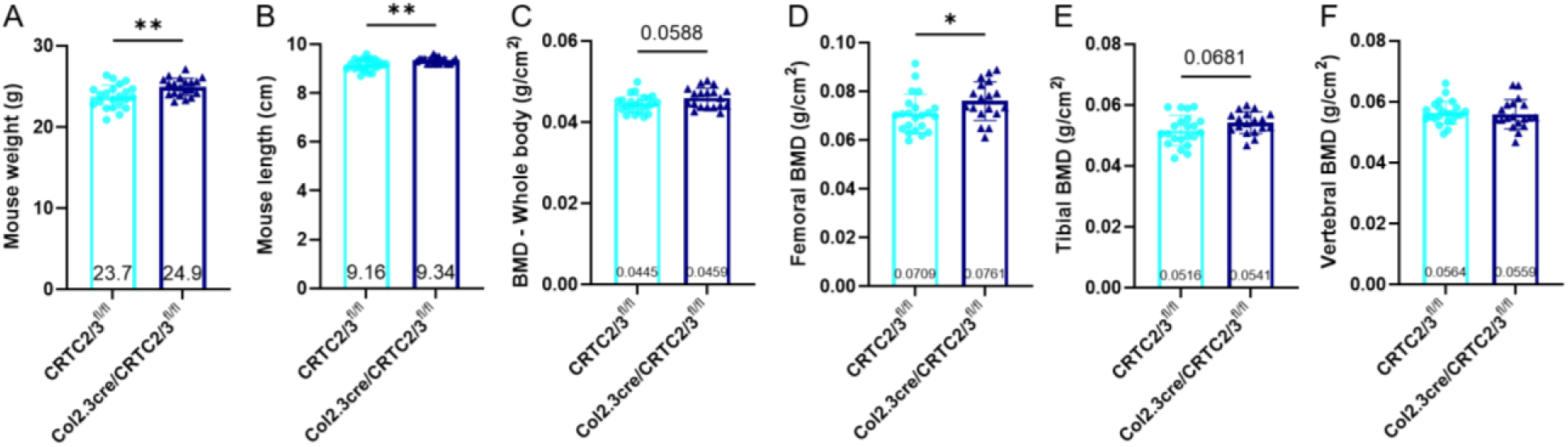
Bone mineral density in young male mice lacking CRTC2 and CRTC3 in osteoblast. (A) Male mouse body weight at euthanasia at 2 months old. (B) Male mouse body length at euthanasia. (C–F) DEXA-PIXImus analysis at euthanasia at 2 months of age to measure bone mineral density (BMD) of (C) whole body (D) femurs, (E) tibiae and (F) vertebrae. Twenty to 23 mice per group. Results are means ± SD. Normality was checked using Anderson-Darling, D’Agostino-Pearson, Shapiro-Wilk and Kolmogorov-Smirnov tests. If data followed a normal distribution, a Welch’s t test was used. * shows significance when p<0.05, **p<0.01

Female conditional knockout mice did not show any obvious phenotype with no changes in body weight, length or BMD at any site. Since the female mice showed less of a phenotype than the males, we have included all the data with female mice as Supplemental Data (Supplemental Figures 1-6).

NanoCT (nCT) analyses of the femurs confirmed the bone changes observed by DEXA-PIXImus, in double conditional knock-out male mice. These mice showed increased trabecular bone volume (BV/TV, Figure 2A) with increased trabecular thickness (Tb.Th, Figure 2B), decreased trabecular separation (Tb.Sp, Figure 2C) and increased trabecular number (Tb.N, Figure 2D) suggesting an increase in osteoblast activity and decreased osteoclast activity. In the metaphysis, male *Crtc2/3^ob-/-^* mice showed a non-significant increased cortical thickness (Figure 2E) without any change in cortical porosity (Figure 2F) but higher cortical bone mineral density (Cortical BMD, Figure 2G). The same phenotype is observed in the diaphysis with a non-significant increase in the cortical thickness (Figure 2H), no changes in cortical porosity (Figure 2I) and bone mineral density (Figure 2J). As with the DEXA-PIXImus analyses, nCT confirmed that CRTC2 and CRTC3 deletion did not affect bone microarchitecture in females with no changes in trabecular bone volume, separation or number (Supplemental Figure 1), although double conditional KO female mice had decreased trabecular thickness (Supplemental Figure 1H). This sexual dimorphism is also seen in the cortical bone in the metaphysis and the diaphysis with no changes in cortical thickness, porosity or bone mineral density (Supplemental Figure 1).

**Figure 2:**
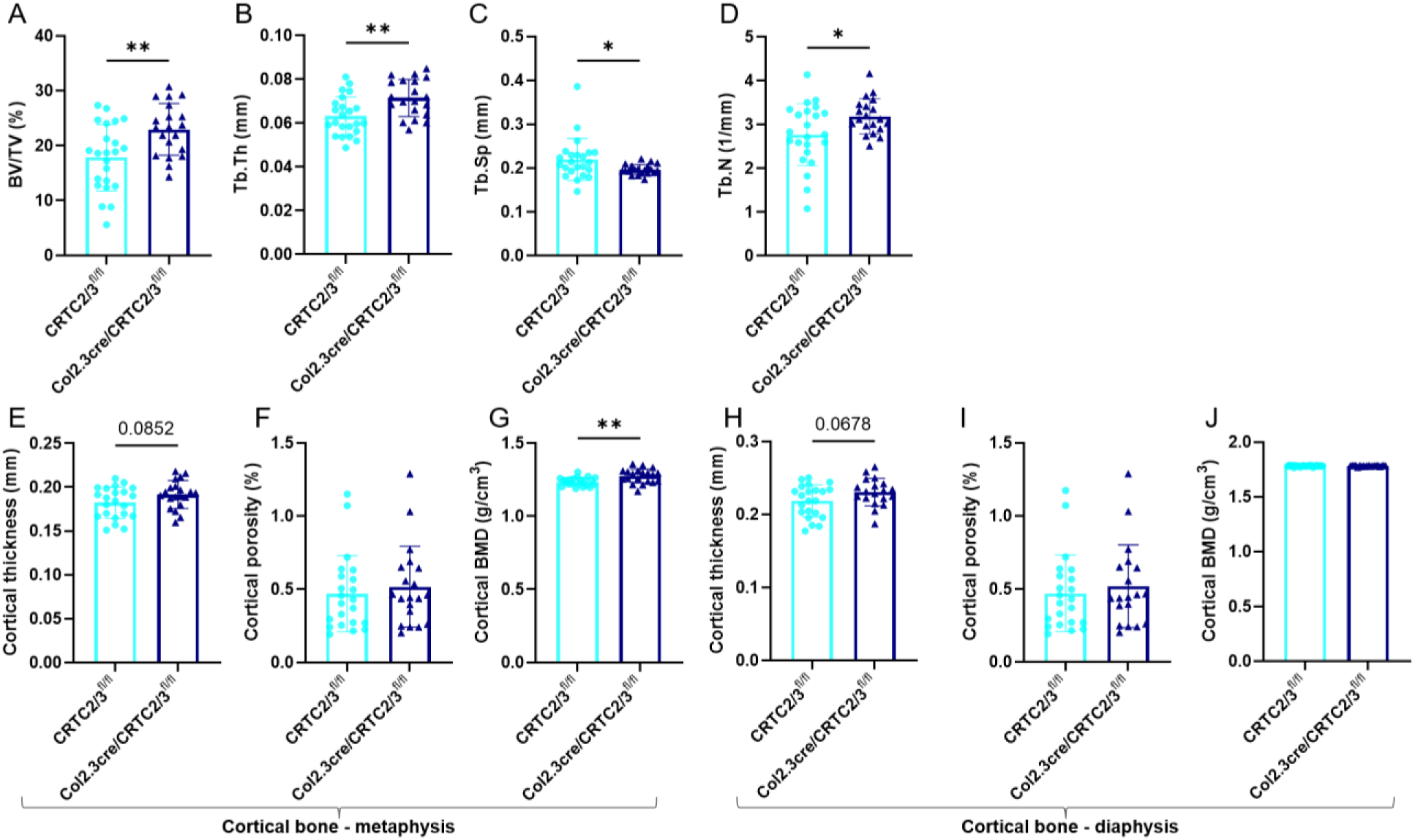
Bone microarchitecture in young male mice lacking CRTC2 and CRTC3 in the osteoblast. Femurs of 2-month-old male mice were prepared for high-resolution nCT and images were reconstructed with NRecon software. (A–E) A 2700 μm volume corresponding to 300 slices of the mid-metaphysis was examined for trabecular bone microarchitecture: (A) trabecular bone volume (BV/TV), (B) trabecular thickness (Tb.Th), (C) trabecular separation (Tb.Sp), and (D) trabecular number (Tb.N). (F-G) A 2250 μm cortical volume corresponding to 250 slices of the metaphysis was examined for (E) cortical thickness, (F) cortical porosity and (G) cortical bone mineral density. (H-J) A 900 μm cortical volume corresponding to 100 slices of the mid-diaphysis was examined for (H) cortical thickness, (I) cortical porosity and (J) cortical bone mineral density. Twenty-one to 22 mice per group; results are means +/- SD. Normality was checked using Anderson-Darling, D’Agostino-Pearson, Shapiro-Wilk and Kolmogorov-Smirnov tests. If data follow a normal distribution, a Welch’s t test was used. If data do not follow a normal distribution (C, F, I), a Mann-Whitney test was used. * shows significance when p<0.05, **p<0.01.

**Supplemental Figure 1:**
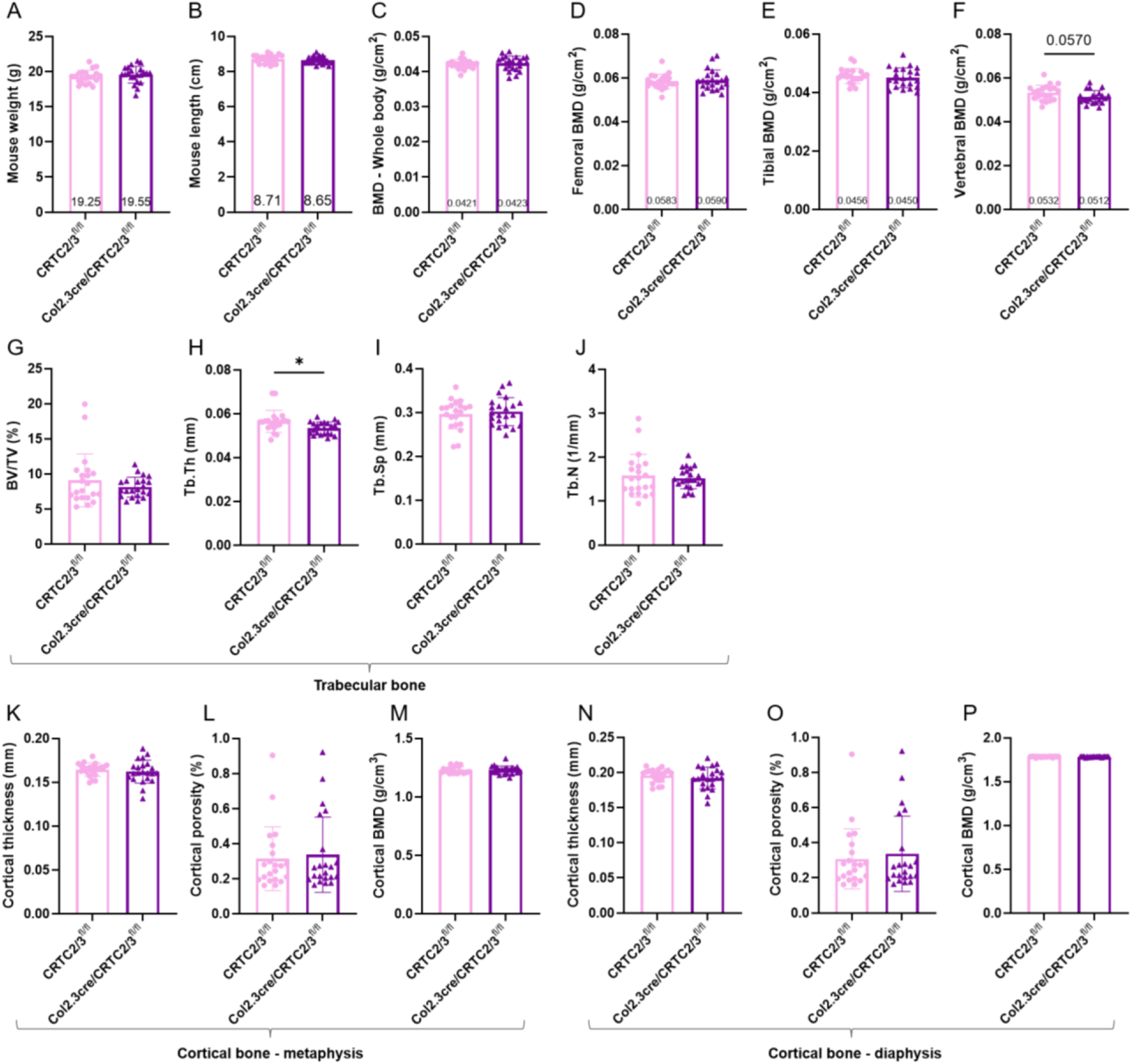
Bone mineral density and bone microarchitecture in young female mice lacking CRTC2 and CRTC3 in osteoblast. (A) Female mouse body weight at euthanasia at 2 months old. (B) Female mouse body length at euthanasia. (C-F) DEXA-PIXImus analysis at euthanasia at 2 months of age to measure bone mineral density (BMD) of (C) whole body (D) femurs, (E) tibiae and (F) vertebrae. (G-P) Femurs of female mice at 2 months old were prepared for high-resolution nCT and a 2700 pm volume was examined for trabecular bone microarchitecture: (G) trabecular bone volume (BV/TV), (H) trabecular thickness (Tb.Th), (I) trabecular separation (Tb.Sp), (J) trabecular number (Tb.N). (K-M) A 2250 μm cortical volume within the metaphysis was examined for (K) cortical thickness, (L) cortical porosity and (M) cortical bone mineral density. (N-P) A 900 μm cortical volume in the mid-diaphysis was examined for (N) cortical thickness, (O) cortical porosity and (P) cortical bone mineral density. Twenty to 23 mice per group; results are means +/- SD. Normality was checked using Anderson-Darling, D’Agostino-Pearson, Shapiro-Wilk and Kolmogorov-Smirnov tests. If data follow a normal distribution, a Welch’s t test was used. If data do not follow a normal distribution (E, G, H, I, J, L, O), a Mann-Whitney test was used. * shows significance when p<0.05

### Male adult mice with osteoblastic deletion of CRTC2/3 showed a slight bone phenotype

At 6 months-old, at euthanasia, male *Crtc2/3^ob-/-^* mice had a slight but not significant increase in body weight (Figure 3A) and no differences in body length (Figure 3B). Unlike the 2-month-old male mice, they did not have increased BMD in the whole body (Figure 3C), femurs (Figure 3D), tibiae (Figure 3E) or vertebrae (Figure 3F). Similar to the observations at 2 months of age, adult female double conditional KO mice did not show any modifications in body weight, length or BMD at any sites (Supplemental Figure 2).

**Figure 3:**
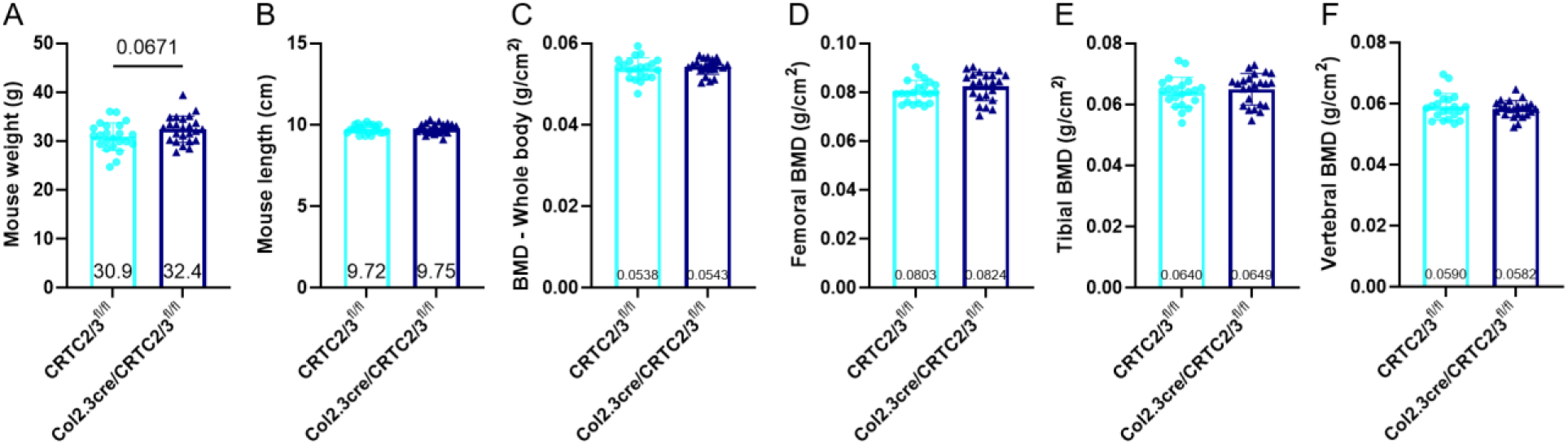
Bone mineral density in adult male mice with deletion of CRTC2 and CRTC3 in the osteoblast. (A) Male mouse body weight at euthanasia at 6 months old. (B) Male mouse body length at euthanasia. (C–F) DEXA-PIXImus analysis at euthanasia at 6 months of age in male mice to measure bone mineral density (BMD) of (C) whole body (D) femurs, (E) tibiae and (F) vertebrae. Twenty-one to 23 mice per group. Results are means ± SD. Normality was checked using Anderson-Darling, D’Agostino-Pearson, Shapiro-Wilk and Kolmogorov-Smirnov tests. If data follow a normal distribution, a Welch’s t test was used. If data do not follow a normal distribution (B, F), a Mann-Whitney test was used.

**Supplemental Figure 2:**
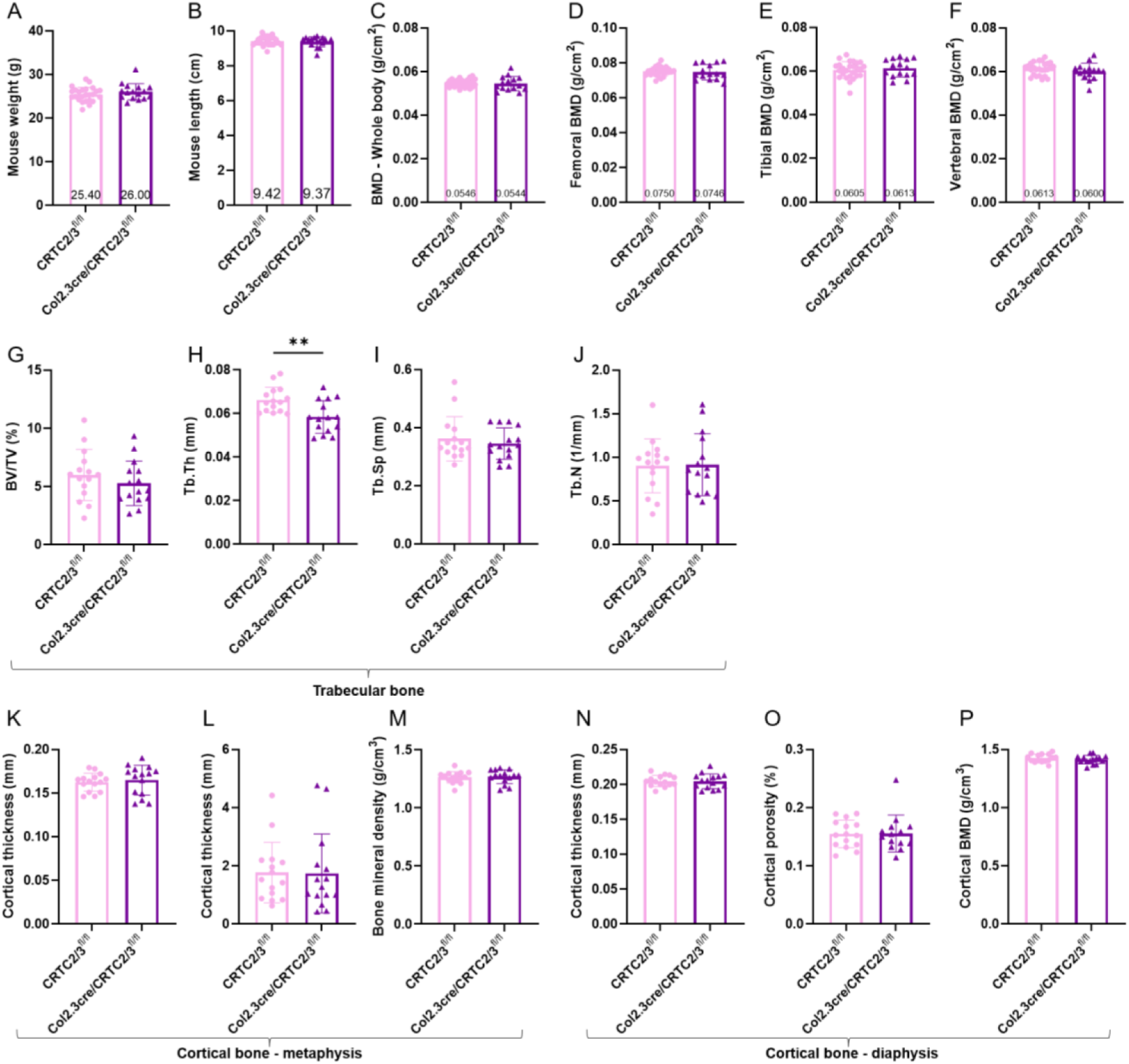
Bone mineral density and bone microarchitecture in adult female mice with deletion of CRTC2 and CRTC3 in the osteoblast. (A) Female mouse body weight at euthanasia at 6 months old. (B) Female mouse body length at euthanasia. (C-F) DEXA-PIXImus analysis at euthanasia at 6 months of age to measure bone mineral density (BMD) of (C) whole body (D) femurs, (E) tibiae and (F) vertebrae. (G-P) Femurs of female mice at 6 months old were prepared for high-resolution nCT and a 2700 μm volume was examined for trabecular bone microarchitecture: (G) trabecular bone volume (BV/TV), (H) trabecular thickness (Tb.Th), (I) trabecular separation (Tb.Sp), (J) trabecular number (Tb.N). (K-M, A 2250 μm cortical volume within the metaphysis was examined for (K) cortical thickness, (L) cortical porosity and (M) cortical bone mineral density. (N-P) A 900 μm cortical volume in the mid-diaphysis was examined for (N) cortical thickness, (O) cortical porosity and (P) cortical bone mineral density. Fifteen to 26 mice per group; results are means +/- SD. Normality was checked using Anderson-Darling, D’Agostino-Pearson, Shapiro-Wilk and Kolmogorov-Smirnov tests. If data follow a normal distribution, a Welch’s t test was used. If data do not follow a normal distribution (B, I, L, P), a Mann-Whitney test was used. * shows significance when p<0.05

While double conditional KO adult male mice did not have higher trabecular bone volume (BV/TV, Figure 4A) which had been observed in males at 2 months of age, they showed a non-significant increase in trabecular thickness (Tb.Th, Figure 4B) while having a significant increase in trabecular separation (Tb.Sp, Figure 4C) suggesting higher osteoblast and osteoclast activities in the adult conditional knockout male mice. In female mice, similar to 2-month-old female mice, deletion of CRTC2 and CRTC3 did not affect the trabecular bone volume, but unexpectedly, the double conditional KO female mice at 6 months of age showed decreased trabecular thickness (Tb.Th, Supplemental Figure 2H) without any changes in trabecular separation (Tb.Sp, Supplemental Figure 2I) or number (Tb.N, Supplemental Figure 2J), suggesting decreased osteoblast activity. Similar to 2-month-old mice, 6-month-old double conditional KO females did not show any changes in cortical bone at either the metaphysis or diaphysis (Supplemental Figure 2K-P).

**Figure 4:**
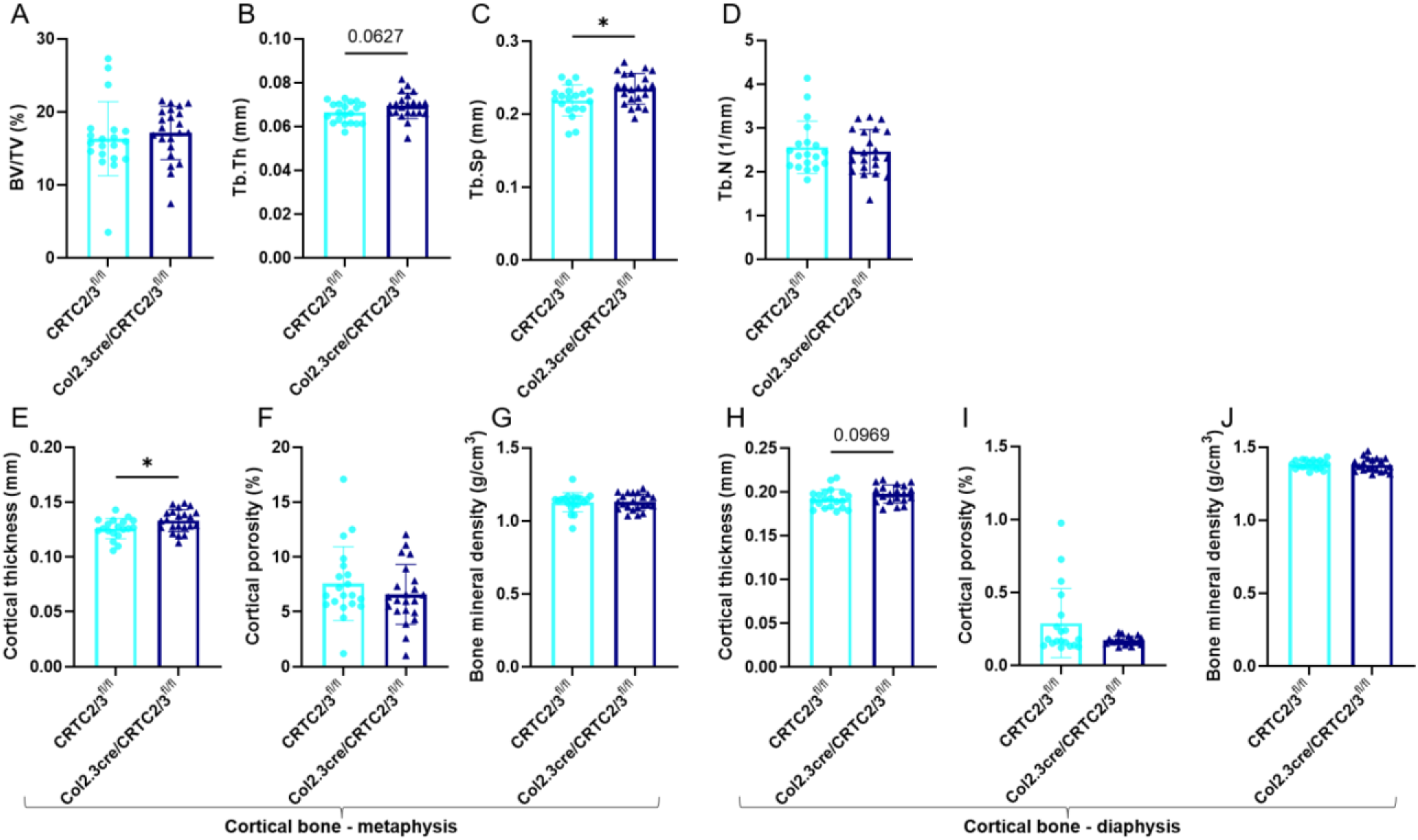
Bone microarchitecture in adult male mice with deletion of CRTC2 and CRTC3 in the osteoblast. Femurs of 6-month-old male and female mice were prepared for high-resolution nCT and images were reconstructed with NRecon software. A 2700 μm volume corresponding to 300 slices of the mid-metaphysis was examined for trabecular bone microarchitecture. A 2250 μm cortical volume corresponding to 250 slices of the metaphysis was examined for bone microarchitecture. A 900 μm cortical volume corresponding to 100 slices of the mid-diaphysis was examined for cortical bone microarchitecture. (A) trabecular bone volume (BV/TV), (B) trabecular thickness (Tb.Th), (C) trabecular separation (Tb.Sp), and (D) trabecular number (Tb.N). (E-G) Metaphyseal cortical bone architecture: (E) cortical thickness, (F) cortical porosity and G) cortical bone mineral density. (H-J) Mid-diaphysis cortical bone architecture: (H) cortical thickness, (I) cortical porosity and (J) cortical bone mineral density. Twenty mice per group; results are means +/- SD. Normality was checked using Anderson-Darling, D’Agostino-Pearson, Shapiro-Wilk and Kolmogorov-Smirnov tests. If data follow a normal distribution, a Welch’s t test was used. If data do not follow a normal distribution (D, K), a Mann-Whitney test was used. * shows significance when p<0.05

### Deletion of CRTC2 and CRTC3 in the osteoblast caused a change in trabecular gene expression in young male mice

To better understand the effect of osteoblastic deletion of *Crtc2* and *Crtc3* in young mice, and since these are transcriptional co-activators, we extracted RNA from the trabecular and cortical areas of tibiae and conducted qPCR analyses. In the trabecular area of young male mice, deletion of *Crtc2* and *Crtc3* caused a decrease in *Runx2* (Figure 5A), *Sost* (Figure 5E) and *Rankl* (Figure 5G) expression, no changes in *Alpl* gene expression (Figure 5C) and increased *Col1α1, Ibsp*, and *Bglap* gene expression (Figure 5B, 5D and 5F). There were no significant changes in gene expression in the cortical bone. As with the bone analyses, the female mice showed very few changes in gene expression, although these mimicked those of the males, but were lesser, with a trend to an increase in *Col1a1* in trabecular bone, and a significant decrease in *Sost* expression in cortical bone (Supplemental Figure 3).

**Figure 5:**
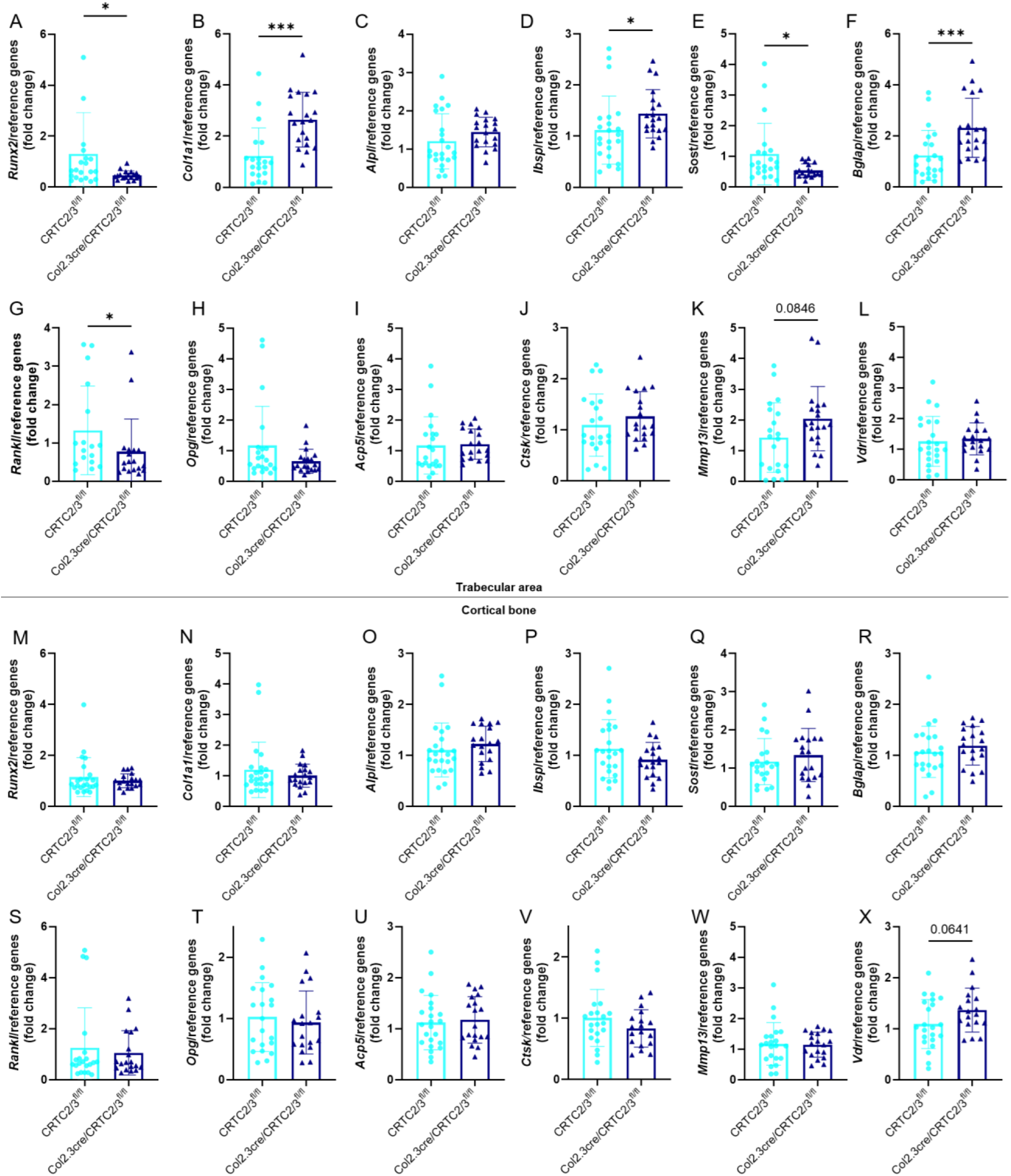
Osteoblastic gene expression in young male mice with deletion of CRTC2 and CRTC3 in the osteoblast. Two tibiae from each animal, at 2 months of age, were divided into subcortical trabecular-rich bone, bone marrow, and cortical bone (osteocyte-rich bone). Total RNA was isolated and qRT-PCR was performed. Osteoblastic gene expression was measured in (A–L) trabecular-rich tibial bone in male mice: (A) *Runx2*, (B) *Type 1 collagen* (*Col1a1*), (C) *Alkaline Phosphatase (Alpl)*, (D) *Bone sialoprotein* (*Ibsp*), (E) *Sost*, (F) *Osteocalcin* (*Bglap*), (G) *Rankl,* (H) *Opg,* (I) *Acp5,* (J) *Ctsk,* (K) *Mmp13,* and (L) *Vdr;* and (M–X) osteocyte-rich cortical tibial bone in male mice: (M) *Runx2*, (N) *Col1a1*, (O) *Alpl,* (P) *Ibsp*, (Q) *Sost*, (R) *Bglap*, (S) *Rankl*, (T) *Opg*, (U) *Acp5*, (V) *Ctsk*, (W) *Mmp13*, and (X) *Vdr*. Normality was checked using Anderson-Darling, D’Agostino-Pearson, Shapiro-Wilk and Kolmogorov-Smirnov tests. If data followed a normal distribution, a Welch’s t test was used. If data did not follow a normal distribution (A, B, D, E, F, G, H, I, J, K, M, N, O, P, Q, R, S, T, V, W, X), a Mann-Whitney test was used. * shows significance when p<0.05, ***p<0.001.

**Supplemental Figure 3:**
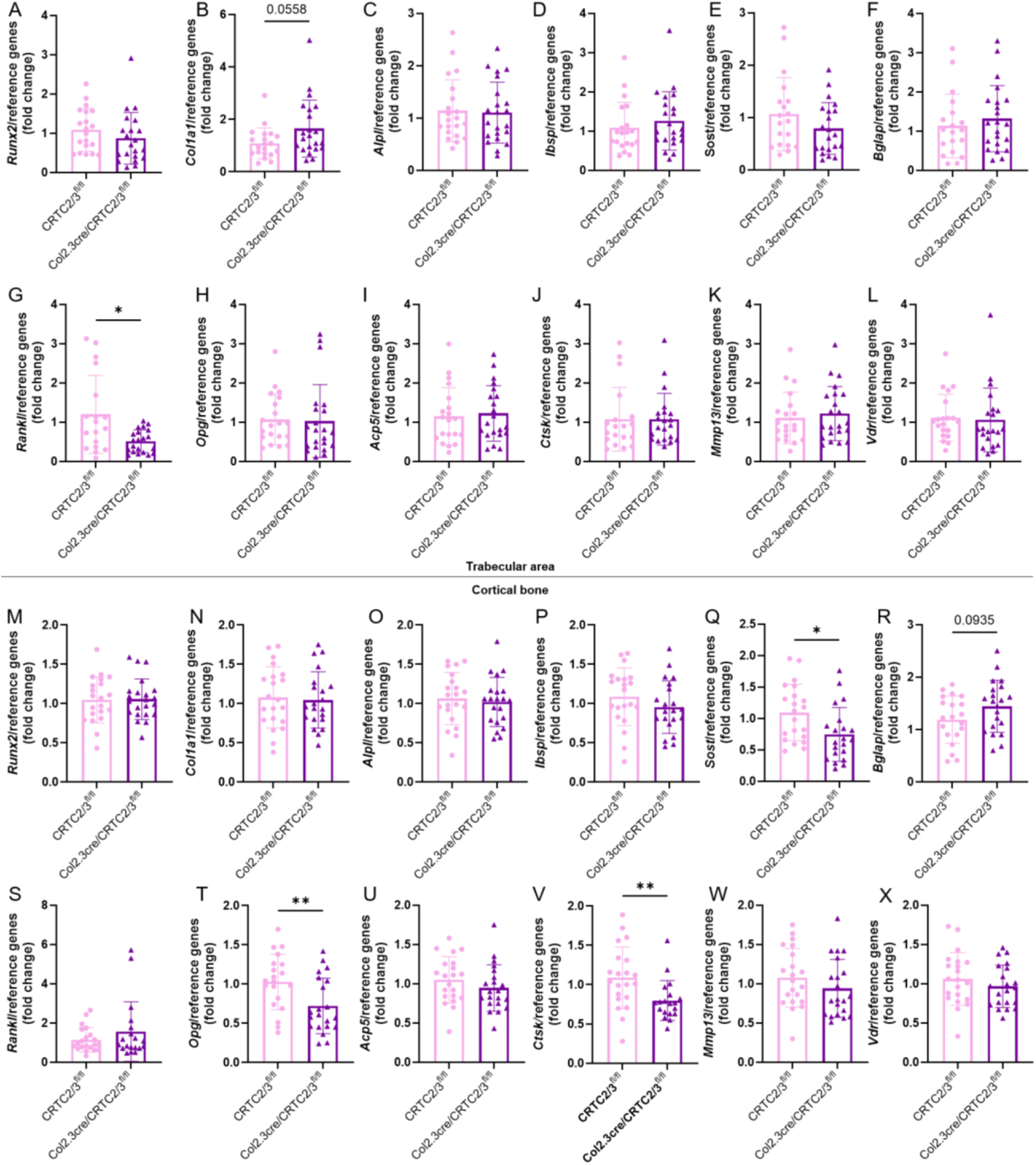
Osteoblastic gene expression in young female mice with deletion of CRTC2 and CRTC3 in the osteoblast. Two tibiae from each female animal, at 2 months of age, were divided into subcortical trabecular-rich bone, bone marrow, and cortical bone (osteocyte-rich bone).), Total RNA was isolated and qRT-PCR was performed. Osteoblastic gene expression was measured in (A-L) trabecular-rich tibial bone: (A) *Runx2,* (B) *Type 1 coflagen (Co/1a1),* (C) *Alkaline Phosphatase (Alpl),* (D) *Bone Sialoprotein (Ibsp),* (E) Sosf, (F) *Osteocalcin (Bglap),* (G) *Rankl,* (H) *Opg,* (I) *Acp5,* (J) *Ctsk,* (K) *Mmpl3,* and (L) *Vdr*; and (M-X) osteocyte-rich cortical tibial bone: (M) *Runx2,* (N) *Collal*, (O) *Alpl,* (P) *Ibsp,* (Q) Sosf, (R) *Bglap,* (S) *Rankl,* (T) *Opg,* (U) *Acp5,* (V) *Ctsk,* (W) *Mmp13,* and (X) *Vdr.* Eighteen to 23 mice per group; results are means +/- SD. Normality was checked using Anderson-Darling, D’Agostino-Pearson, Shapiro-Wilk and Kolmogorov-Smirnov tests. If data followed a normal distribution, a Welch’s t test was used. If data did not follow a normal distribution (A, B, C, D, E, F, G, H, I, J, K, L, Q, W), a Mann-Whitney test was used. * shows significance when p<0.05, **p<0.01.

### Deletion of CRTC2 and CRTC3 in the osteoblast caused a decrease in trabecular *Rankl* gene expression

Investigation of expression of genes involved with bone and matrix breakdown, showed that the expression of trabecular *Rankl* was decreased in both male and female 2-month-old mice (Figure 5G and Supplemental Figure 3G). Females at this age also expressed less cortical *Opg* and *Ctsk* (Supplemental Figure 3T and V). *Rankl* transcription *in vitro* is directly regulated by CRTC2 and CRTC3 as we and others have shown^(3, 5-7)^. However, there were no changes in *Rankl* expression in cortical bone at this age.

### Wnt pathway genes are changed by deletion of CRTC2 and CRTC3 in the osteoblast

Since we had found that several *Wnt* genes were increased by PTH in differentiated mouse calvarial cells and these increases seem to be regulated through the SIK/CRTC pathway^(7)^, we investigated the expression of a number of Wnt pathway genes in the mice with osteoblastic deletion of CRTC2/3. We had already found decreases in *Sost* expression which could lead to an increase in the activity of the Wnt pathway and further downstream effects on bone phenotype. In fact, we did find decreases in *Lef1* and *Wnt11,* an increase in *Wisp* and *Tcf7* in males, and decreases in *Dvl3*, *Sfrp4* and *Dkk1* in females (Supplemental Figure 4).

**Supplemental Figure 4:**
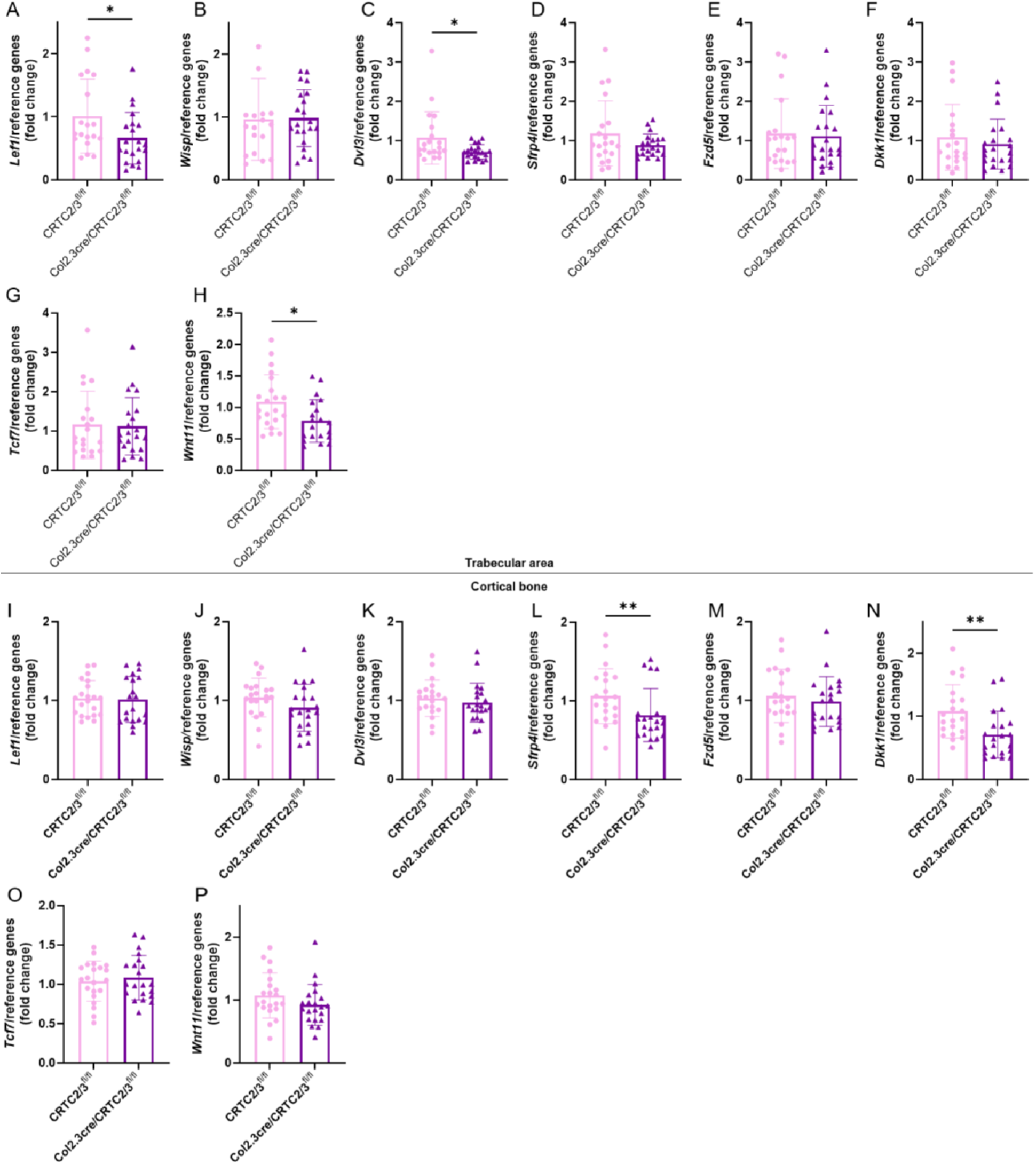
Wnt pathway gene expression in young female mice with deletion of CRTC2 and CRTC3 in the osteoblast. Two tibiae from each female mouse, at 2 months of age, were divided into subcortical trabecular-rich bone, bone marrow, and cortical bone {osteocyte-rich bone}. Total RNA was isolated and qRT-PCR was performed. Osteoblastic gene expression was measured in (A-H) trabecular-rich tibial bone: (A) *Lef1*, (B) Wsp, (C) *Dvl3,* (D) *Sfrp4,* (E) *Fzd5,* (F) *Dkk1,* (G) *Tcf7* and (H} Wnff *1*; and (l-P) osteocyte-rich cortical tibial bone: (I) *Lefl,* (J) *Wisp,* (K) *Dvl3,* (L) *Sfrp4,* (M) *Fzd5,* (N) *Dkk1,* (O) *Tcf7* and (P) *Wnt11.* Eighteen to 23 mice per group; results are means +/- SD. Eighteen to 23 mice per group; Normality was checked using Anderson-Darling, D’Agostino-Pearson, Shapiro-Wilk and Kolmogorov-Smirnov tests. If data followed a normal distribution, a Welch’s I test was used. If data did not follow a normal distribution {A, B, C, D, E, F, G, H I, K, L, N, P}, a Mann-Whitney test was used. * shows significance when p<0.05, **p<0.01.

### Deletion of CRTC2 and CRTC3 in the osteoblast caused a change in trabecular gene expression in adult male mice

Similar to the data with 2-month-old mice, adult male mice showed decreases in *Runx2* and *Sost* expression in trabecular bone (Figure 7A and 7E), but as well, a decrease in *Runx2* and *Alpl* in cortical bone (Figure 6G and 6I). Females also showed a decrease in *Sost* expression in trabecular bone at this age (Supplemental Figure 5). However, there were no significant changes in *Rankl* expression, although an increase in trabecular *Ctsk* expression in males. There were many changes in Wnt pathway gene expression, particularly in the males, increases in *Lef1, Dkk1* and *Wnt11* in trabecular bone (Figure 8); in cortical bone increased *Lef1*, but decreased *Wisp*, *Dvl3*, *Fzd5*, *Dkk1*, *Tcf7* and *Wnt11* (Figure 8). There were fewer changes in the females (Supplemental Figure 6).

**Figure 6:**
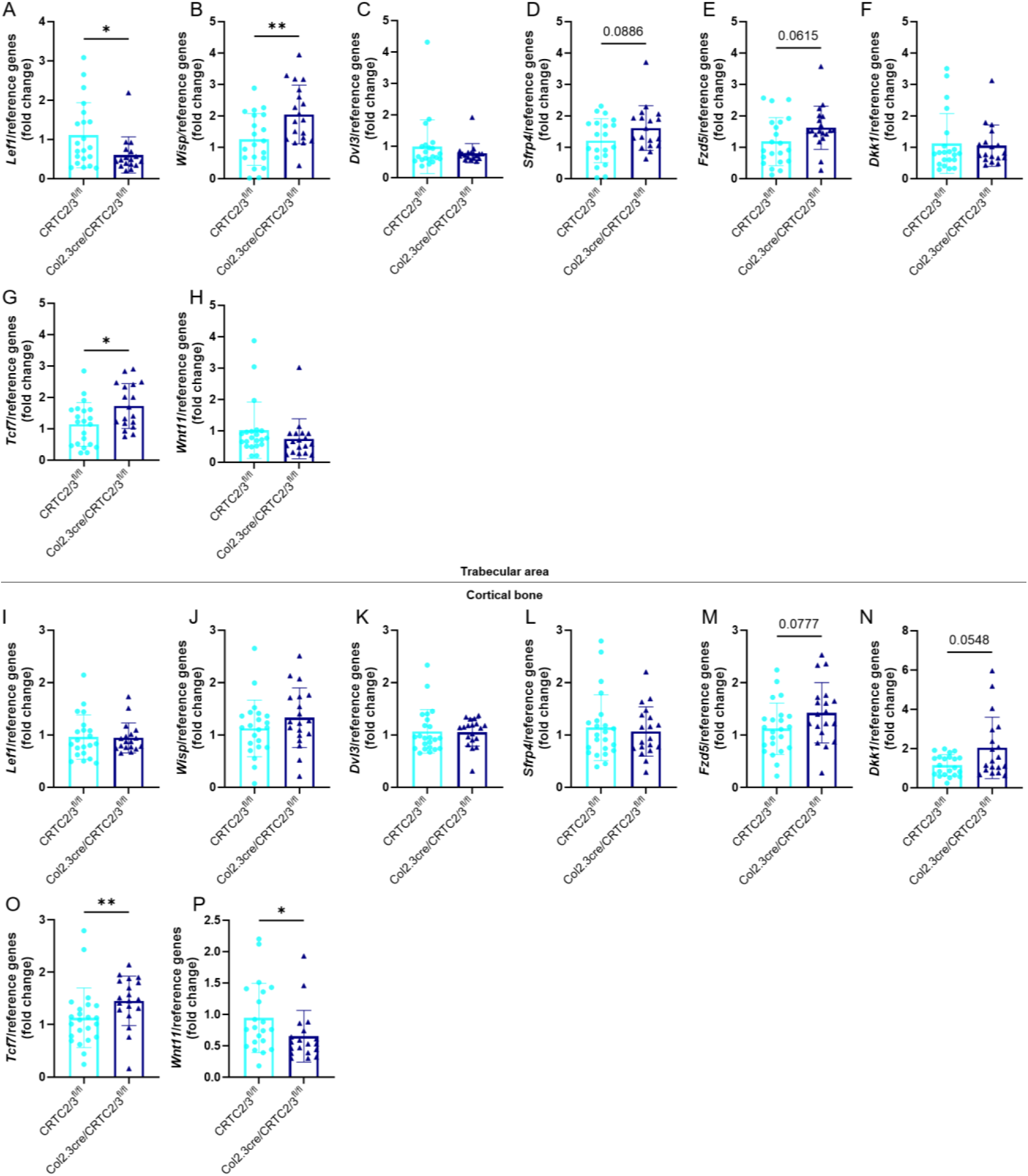
Wnt pathway gene expression in young male mice with deletion of CRTC2 and CRTC3 in the osteoblast. Two tibiae from each animal, at 2 months of age, were divided into subcortical trabecular-rich bone, bone marrow, and cortical bone (osteocyte-rich bone). Total RNA was isolated and qRT-PCR was performed. Osteoblastic gene expression was measured in (A–H) trabecular-rich tibial bone in male mice: (A) *Lef1*, (B) *Wisp*, (C) *Dvl3*, (D) *Sfrp4*, (E) *Fzd5*, (F) *Dkk1*, (G) *Tcf7*, and (H) *Wnt11*; and (I–P) osteocyte-rich cortical tibial bone in male mice: (I) *Lef1*, (J) *Wisp*, (K) *Dvl3*, (L) *Sfrp4*, (M) *Fzd5*, (N) *Dkk1*, (O) *Tcf7*, and (P) *Wnt11*. Eighteen to 23 mice per group; results are means +/- SD. Normality was checked using Anderson-Darling, D’Agostino-Pearson, Shapiro-Wilk and Kolmogorov-Smirnov tests. If data followed a normal distribution, a Welch’s t test was used. If data did not follow a normal distribution (A, C, F, H, I, K, L, N, P), a Mann-Whitney test was used. * shows significance when p<0.05, **p<0.01.

**Figure 7:**
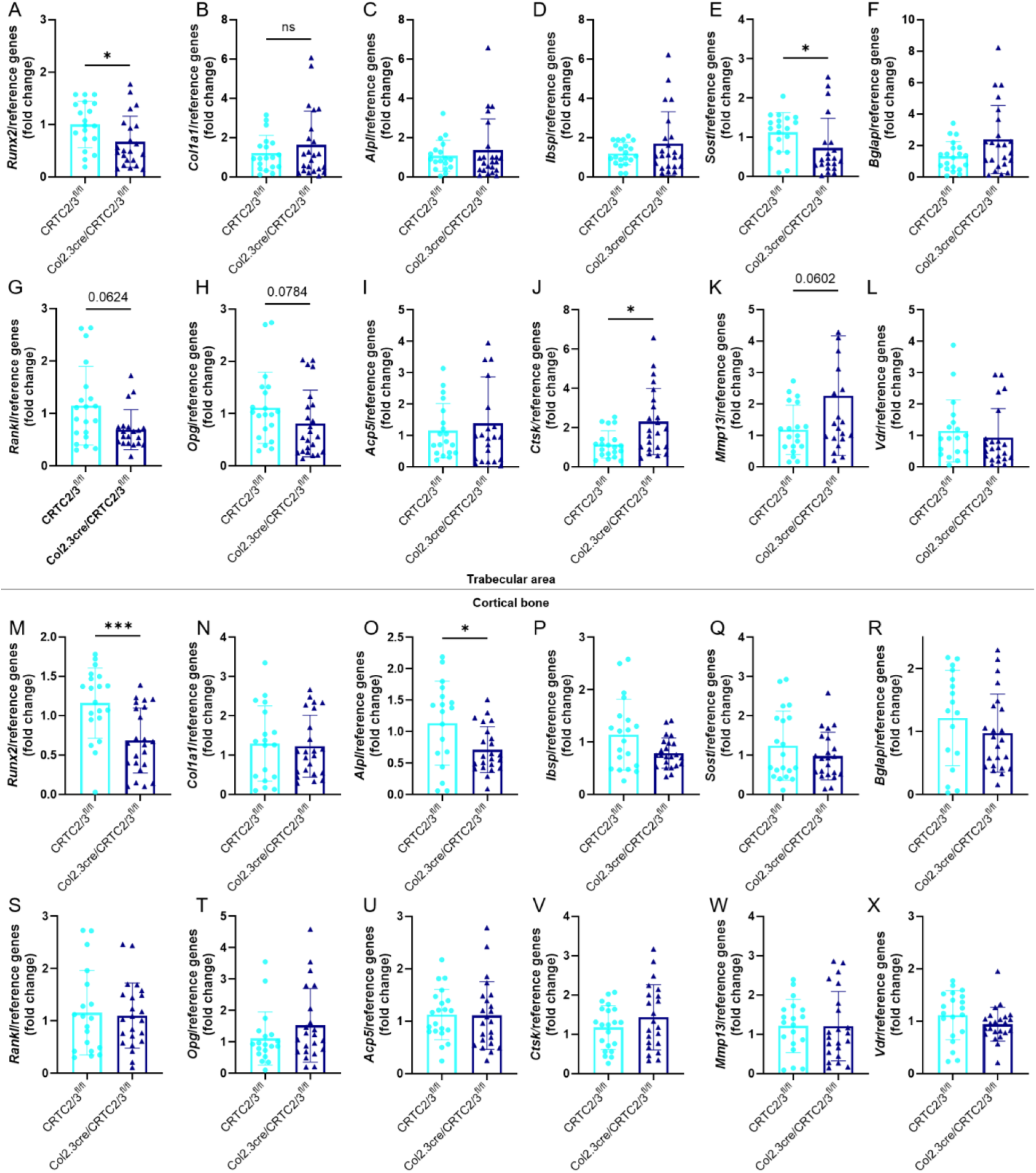
Osteoblastic gene expression in adult male mice with deletion of CRTC2 and CRTC3 in the osteoblast. Two tibiae from each animal, at 6 months of age, were divided into subcortical trabecular-rich bone, bone marrow, and cortical bone (osteocyte-rich bone). Total RNA was isolated and qRT-PCR was performed. Osteoblastic gene expression was measured in (A–F) trabecular-rich tibial bone in male mice: (A) *Runx2*, (B) *Type 1 collagen* (*Col1a1*), (C) *Alkaline Phosphatase (Alpl)*, (D) *Bone sialoprotein* (*Ibsp*), (E) *Sost*, (F) *Osteocalcin* (*Bglap*), (G) *Rankl,* (H) *Opg,* (I) *Acp5,* (J) *Ctsk,* (K) *Mmp13,* and (L) *Vdr*; and (M–X) osteocyte-rich cortical tibial bone in male mice: (M) *Runx2*, (N) *Col1a1*, (O) *Alpl,* (P) *Ibsp*, (Q) *Sost*, (R) *Bglap*, (S) *Rankl*, (T) *Opg*, (U) *Acp5*, (V) *Ctsk*, (W) *Mmp13*, and (X) *Vdr*. Fifteen to 23 mice per group; results are means +/- SD. Normality was checked using Anderson-Darling, D’Agostino-Pearson, Shapiro-Wilk and Kolmogorov-Smirnov tests. If data followed a normal distribution, a Welch’s t test was used. If data did not follow a normal distribution (B, C, D, E, F, G, H, I, J, K, L, N, P, Q, R, S, T, U) a Mann-Whitney test was used. * shows significance when p<0.05, **p<0.01.

**Figure 8:**
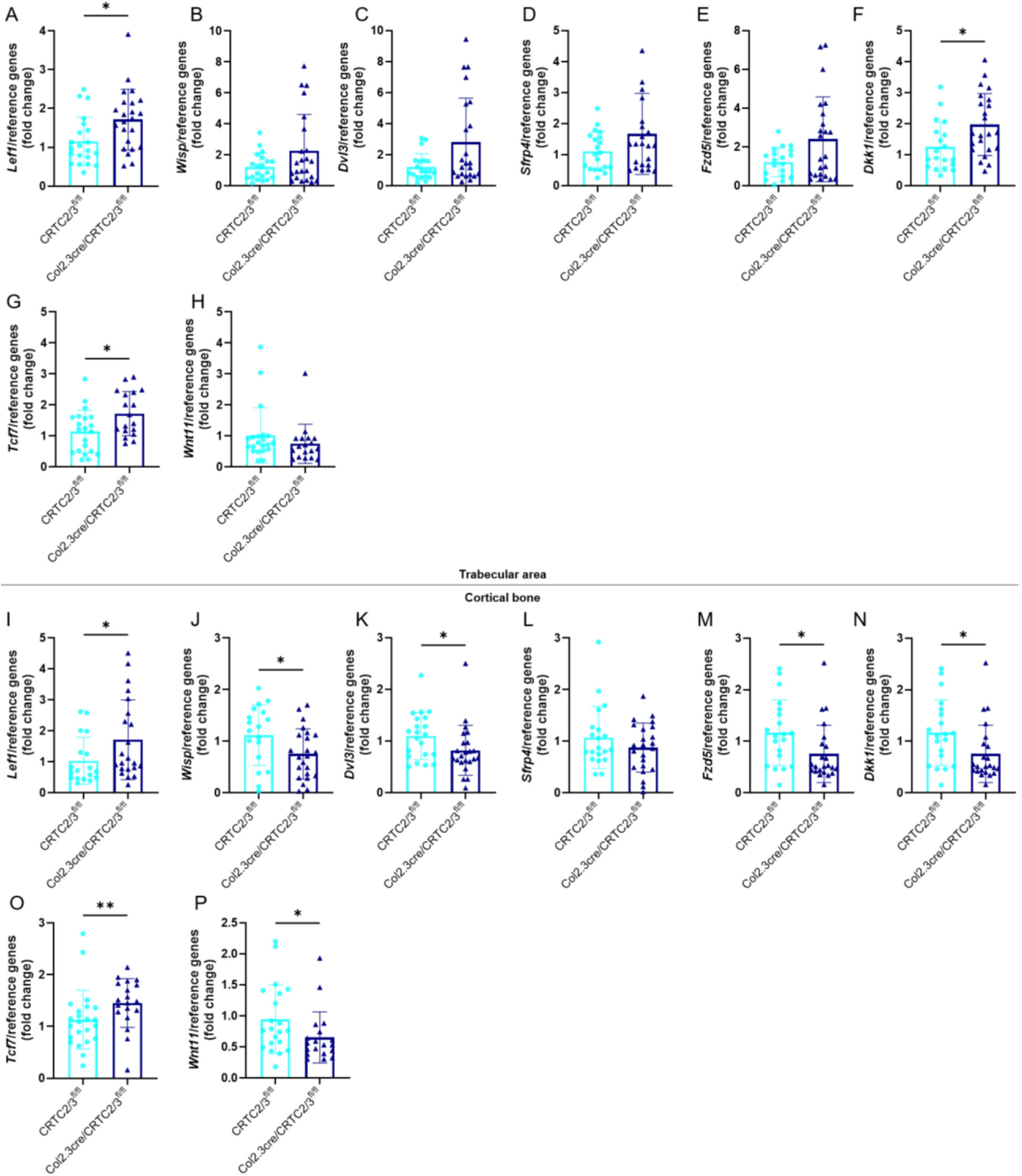
Wnt pathway gene expression in adult mice with deletion of CRTC2 and CRTC3 in the osteoblast. Two tibiae from each animal, at 6 months of age, were divided into subcortical trabecular-rich bone, bone marrow, and cortical bone (osteocyte-rich bone). Total RNA was isolated and qRT-PCR was performed. Osteoblastic gene expression was measured in (A–F) trabecular-rich tibial bone in male mice: (A) *Lef1*, (B) *Wisp*, (C) *Dvl3*, (D) *Sfrp4*, (E) *Fzd5*, (F) *Dkk1*, (G) *Tcf7*, and (H) *Wnt11*; and (I–P) osteocyte-rich cortical tibial bone in male mice: (I) *Lef1*, (J) *Wisp*, (K) *Dvl3*, (L) *Sfrp4*, (M) *Fzd5*, (N) *Dkk1*, (O) *Tcf7*, and (P) *Wnt11*. Fifteen to 23 mice per group; results are means +/- SD. Normality was checked using Anderson-Darling, D’Agostino-Pearson, Shapiro-Wilk and Kolmogorov-Smirnov tests. If data followed a normal distribution, a Welch’s t test was used. If data did not follow a normal distribution (A, B, C, D, E, H, I, K, L, M, N, O), a Mann-Whitney test was used. * shows significance when p<0.05.

**Supplemental Figure 5:**
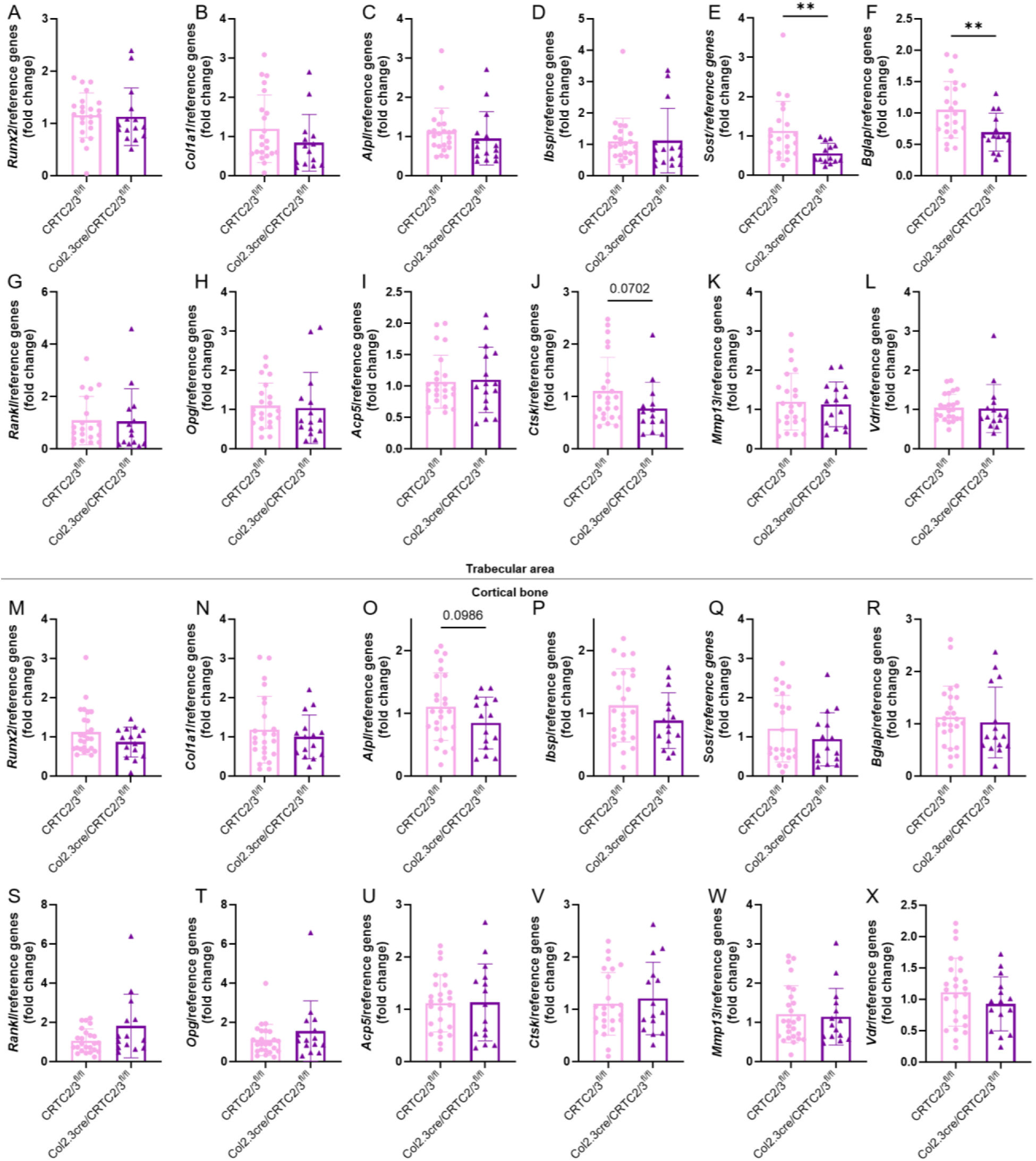
Osteoblastic gene expression in adult female mice with deletion of CRTC2 and CRTC3 in the osteoblast. Two tibiae from each female animal, at 6 months of age. were divided into subcortical trabecular-rich bone, bone marrow, and cortical bone (osteocyte-rich bone). Total RNA was isolated and qRT-PCR was performed. Osteoblastic gene expression was measured in (A-L) trabecular-rich tibial bone; (A) Rvnx2, (B) *Type 1 collagen (Coital),* (C) *Alkaline Phosphatase (A/pl),* (D) *Bone Sialoprotein (Jbsp),* (E) Sosf, (F) *Osteocalcin (Bglap),* (G) *Rankl,* (H) Opg, (I) *Acp5,* (J) *Ctsk,* (K) *Mmpl3,* and (L) *Vdr*; and (M-X) osteocyte-rich cortical tibial bone: (M) *Runx2,* (N) *Col1a1,* (O) *Alpl,* (P) *Ibsp,* (Q) Sosf, (R) *Bglap,* (S) *Rankl,* (T) *Opg,* (U) *Acp5,* (V) *Ctsk,* (W) *Mmp13,* and (X) *Vdr.* Fifteen to 23 mice per group; results are means +/- SD. Normality was checked using Anderson-Darling, D’Agostino-Pearson, Shapiro-Wilk and Komogorov-Smimov tests. If data follow a normal distribution, a Welch’s t.test was used. If data do not follow a normal distribution (A, B, C, D, E, G, H, I, J, L, M, N, Q, R, S, T, V, W, X), a Mann-Whitney test was used. * shows significance when p<0. 05, **p<0.01.

**Supplemental Figure 6:**
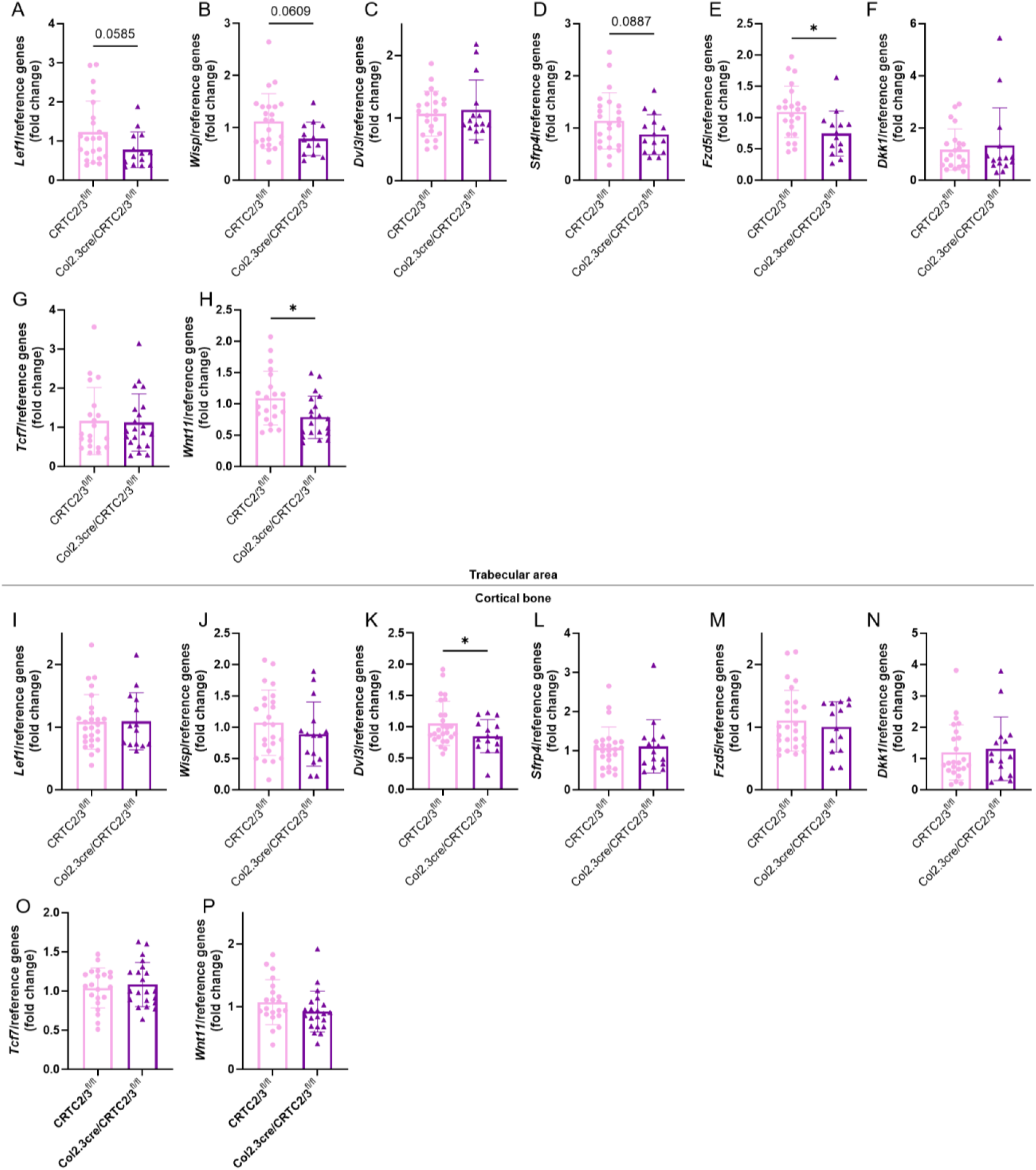
Wnt pathway gene expression in adult female mice with deletion of CRTC2 and CRTC3 in the osteoblast. Two tibiae from each female mouse, at 2 months of age, were divided into subcortical trabecular-rich bone, bone marrow, and cortical bone (osteocyte-rich bone). Total RNA was isolated and qRT-PCR was performed. Osteoblastic gene expression was measured in (A-H) trabecular-rich tibial bone: (A) *Lef1,* (B) W/sp, (C) *Dvl3,* (D) *Sfrp4,* (E) *Fzd5,* (F) *Dkkt,* (G) *Tcf7* and (H) Whff *1*; and (l-P) osteocyte-rich cortical tibial bone: (I) *Lef1,* (J) W/sp, (K) *Dvl3,* (L) *Sfrp4,* (M) *Fzd5,* (N) *Dkk1,* (O) *Tcf7* and (P) *Wnt11.* Fifteen to 23 mice per group; results are means +/- SD. Normality was checked using Anderson-Darling, D’Agostino-Pearson, Shapiro-Wilk and Komogorov-Smirnov tests. If data follow a normal distribution, a Welch’s t test was used. If data do not follow a normal distribution (A, B, C, F, G, H, I, K, N, P), a Mann-Whitney test was used. * shows significance when p<0.05.

## Discussion

The goal of this study was to better understand the role of CRTC2 and CRTC3 *in vivo* in osteoblasts and if its deletion had developmental effects on bone and bone gene expression or decreased gene expression regulated by the PTHR1 pathway. We constitutively deleted CRTC2 and CRTC3, using a *Col1α1* promoter (*Col2.3Cre*)) which targets mature osteoblasts. We found that there were overt effects mainly on male young mice including increased body weight, length and bone mass. These effects declined with age to adulthood. However, we also found a significant decrease in expression of *Runx2*, *Sost* and *Rankl* in the young male mice. The decrease in *Sost* and *Rankl* might cause an increase in growth and bone mass through the Wnt pathway, and through a decrease in osteoclast activity.

We had previously investigated *in vitro* a number of genes which had been shown to be highly regulated by PTH (1-34) in differentiated calvarial osteoblasts^(7)^ and several of whose regulation by PTH had been found to involve the SIK/CRTC pathway. In particular, we determined that *Rankl, Wnt4* and *Wnt7b* required both SIKs and CRTCs for PTH stimulation of their gene expression. Others such as *Sost, Sfrp4* and *Wnt11* were either known or found to be only SIK-dependent, possibly all operating through the alternative arm of SIK action through the Type II HDACs^(4)^, and not through the CRTCs. Thus, it is reassuring that, with deletion of CRTC2 and CRTC3 in osteoblasts *in vivo*, we found a decrease in *Rankl* expression coupled with decreases in *Ctsk*, suggesting less osteoclast activity in these bones. Our observations support the notion that CRTC2 and CRTC3 are major regulators of *Rankl* transcription *in vivo*. It was unexpected that we would find decreases in *Runx2* and *Sost* expression, suggesting that CRTC2 and CRTC3 may have a role in controlling their basal expression. We did not find this for *Sost in vitro*^(7)^. PTH is known to stimulate *Runx2* through the PKA pathway^(13, 14)^ while it decreases *Sost* expression through the PKA/SIK/HDAC4/5 axis^(4)^. It is possible that CRTC2 and CRTC3, bound to bZip transcription factors, act as basal regulators of their transcription. Alternatively, since this was all *in vivo* work, there could be a change in feedback regulation from lesser osteoclast activity and lesser clastokines or lower release of matrix-embedded growth factors such as IGF-1^(15)^ and TGF-β^(16, 17)^, causing a decrease in expression of *Runx2* and *Sost*. We were unable to find changes in *Wnt4* expression, since we found that it is expressed at very low mRNA levels *in vivo*, unlike in calvarial osteoblastic cells *in vitro*.

The sexual dimorphism we observed suggest that either estradiol or the XX chromosomes modulate the effects of CRTC2/3 on gene expression in bones of the female mice. In fact, decreased *Crtc2* mRNA was observed in the paraventricular nucleus neurons of adrenalectomized diestrous and proestrous females vs. males^(18)^, which the authors ascribed to an underlying influence of estradiol. It would be interesting to try the Four Core Genotype mouse model to determine if it is an effect of estradiol or the contributions from the X and Y chromosomes^(19)^.

In summary, we have found moderate effects on the bone phenotype of young male mice with deletion of both CRTC2 and 3 in the osteoblastic lineage, which are reflected in decreases in osteoblastic gene expression, especially *Rankl*. This supports the data from work *in vitro* and forms a basis for investigation of the role of these co-activators in PTH action *in vivo*.

## Materials and methods

### Animals

All mice were on a C57Bl/6J background. The mice were fed with a mouse standard diet (PicoLab® Rodent Diet 20, 5053, LabDiet) containing calcium (0.81%), phosphorus (0.63%) and vitamin D (2.2 IU/g). All mice were kept on a 12 h light/dark cycle with standard rodent chow and water *ad libitum*. All mouse-related experimental procedures were performed in accordance with approved protocols of the Institutional Animal Care and Use Committee of New York University Grossman School of Medicine or Rutgers University. To delete CRTC2 and CRTC3 in osteoblasts, we crossed *Col2.3*Cre mice in several steps with *Crtc2/3^fl/fl^*mice. This generated a colony of breeding pairs of *Col2.3*Cre/*Crtc2/3^fl/fl^*males and *Crtc2/3^fl/fl^* females to generate the numbers of animals required for the experiments. Only male mice were used for the Cre drivers because of the issues raised about transfer of Cre maternally^(20)^. At 2 months of age, animals were examined for body mass, length and assessed by DEXA-PIXImus for bone mineral density (BMD). Prior to being euthanized, young animals were injected with tetracycline (20 mg/kg, Sigma) then calcein (10 mg/kg, Sigma) 4 and 1 days before euthanasia. For the long-term experiment, mice were examined monthly for body mass and bone mineral density (BMD) assessed by DEXA-PIXImus. Adult mice were euthanized at 6 months old. The adult animals received double fluorescent injections (tetracycline and calcein) at 2 different time points (D7 and D2 before death) to measure mineral apposition rate (MAR) and bone formation rate (BFR).

### DEXA-PIXImus Analyses

At euthanasia or 2 and 6 months old, young mice were weighed and a DEXA-PIXImus used to assess changes in post-cranial skeletal areal bone mineral density (BMD) by an independent blinded person. On the day of BMD measurement, the machine is warmed up and the machine is calibrated by measuring the phantom. The program is composed of several impulses of two different X-ray beams: LE (’Low Energy’ X-ray attenuation, voltage at 35 kV, current at 0.5 mA for 15 seconds) and HE (’High Energy’ X-ray attenuation, voltage at 80 kV, current at 0.5 mA for 3 seconds). Then, the mouse is anesthetized with ketamine (100 mg/kg) / xylazine (10 mg/kg) and laid down with the head in the circular groove of the specimen tray. The tail is laid away from the main body to be sure that it does not cross any of the long bones. With gentle traction, the spine is straightened, and the skull is aligned to the sagittal plane. Legs are moved away from the body with the front legs at a 45° angle to the spine to prevent overlap and femurs at a 90° angle to the spine. Every image is checked immediately after each DEXA-PIXImus scan. If the mouse moved, it is rescanned. Accurate measurements were taken, always in the same fashion, by a person blinded to the animal. Bone mineral density (BMD) was measured by defined region of interest of the whole body (excluding the skull and the ear tag), the right femurs (from the hip to the middle of the knee), the right tibiae (the middle of the knee to the middle of the ankle), and lumbar vertebrae (from the ribs to the hips). Absolute BMD values were expressed in g/cm^2^.

### Nano-Computed Tomography (nCT)

After euthanasia, right femurs were dissected, cleaned of soft tissue, and fixed in 70% ethanol for at least 3 weeks and subjected to nCT analyses. The samples were scanned in batches of six at a nominal resolution (pixels) of 9 μm using a high-resolution micro-computed tomography system (nCT, SkyScan 1272, SkyScan, Ltd., Kartuizersweg, Kontich, Belgium). The following imaging parameters were used: 70 kV, 142 μA, and an aluminum 0.5 mm filter. All images were reconstructed using NRecon (Skyscan), a 3D morphometry evaluation program, with the following parameters: beam-hardening correction of 41 and Gaussian smoothing (factor 2). The reconstructed data were binarized using a thresholding of 60–255 for trabecular bone and 80-255 for cortical bone. CTAn software (SkyScan, Kartuizersweg, Kontich, Belgium) was used for all three-dimensional volumetric analyses of trabecular bone and two-dimensional analyses of cortical bone. Bone mineral density of trabecular and cortical bone was determined from the binary data based on a calibration curve of hydroxyapatite standards. The nCT measurements follow the guidelines reported by Bouxsein *et al*.^(21)^. For 2 and 6 month-old mice, a 2700 μm volume corresponding to 300 slices of the metaphysis that began 20 slices below the growth plate was examined for trabecular bone microarchitecture, a 2250 μm cortical volume corresponding to 250 slices of the metaphysis that began 70 slices below the growth plate and a 900 - 1800 μm cortical volume (900 μm for mice at 2 months of age and 1800 μm for 6-month-old mice) corresponding to 100 - 200 slices of the mid-diaphysis were examined for cortical bone microarchitecture. All analyses were done blinded by 2 different persons.

### RNA Isolation and Quantitative RT-PCR Analyses

Both tibiae per animal were dissected and cleared of soft tissues. Then, tibiae were separated into different bone compartments such as distal and proximal ends of the tibiae, corresponding to the subcortical trabecular rich region and the growth plate, was first dissected; bone marrow by centrifuging the bone and the remaining cortical bone containing predominantly osteocytes. Total RNA was extracted using a TRIzol kit (Sigma). cDNA was synthesized from 1 μg of total RNA using TaqMan® Reverse Transcription Reagents (Life Technologies, Inc.). SYBR® Green Master Mix was used for quantitative real-time RT-PCR using a Mastercycler® realplex^2^ instrument (Eppendorf). mRNA expression was calculated using the following formula 2^^-(ΔΔCt)^. The levels of mRNA expression were normalized to geometrical means with *Gapdh, β-actin,* and *Hprt* expression as internal controls and then expressed as fold values compared with the *Crtc2/3^ob+/+^* mice. The qRT-PCR primers are listed in Table 1. All qRT-PCR shown were done on trabecular-rich or cortical-rich regions.

### Statistical Analyses

Values are presented as the means ± S.D. and shown as individual values. Data were analyzed for normal distribution by Anderson-Darling, D’Agostino-Pearson, Shapiro-Wilk and Kolmogorov-Smirnov tests. If normal distribution was not achieved, we used rank transformation and confirmed the normality by the previous tests. The equality of variance was determined with Levene’s test. If both normal distribution and equality of variance were demonstrated we used Student’s t Test using SigmaStat software (SPSS Sciences, Chicago IL) and GraphPad Prism 8 (2019 GraphPad Software, Inc., La Jolla, CA, USA). If equivalence of variance was not achieved, we used Welch’s t Test. If we could not perform a parametric test, we used a non-parametric test such as the Mann-Whitney test.

## Acknowledgements

This work was supported by NIH grants R01 DK047420 and S10 OD010751 (to NCP). The authors have no conflicts of interest.

## Authors contributions

Authors’ roles: Study design: CLH and NCP. Study conduct: CLH. Data collection: CLH, JJ, WP, ZH, AM, SS, and NCP. Data analysis and interpretation: CLH and NCP. Drafting manuscript: CLH and NCP. Revising manuscript content: CLH, WP, JJ and NCP. Approving final version of manuscript: CLH, JJ, WP, ZH, AM, SS, and NCP. CLH takes responsibility for the integrity of data analysis.

Supplemental data have been included with the submission.

